# High-Quality Draft Genome Assemblies and Comparative Genomics of Three *Bouteloua* Species (Poaceae: Chloridoideae)

**DOI:** 10.1101/2025.10.15.682643

**Authors:** Mathavan A. Ganesan, Charles O. Hale, M. Cinta Romay, Edward S. Buckler, Michelle C. Stitzer

## Abstract

*Bouteloua* is a PACMAD grass genus that dominates much of the central and southern North American prairies and plains, where species such as sideoats grama (*B. curtipendula*), blue grama (*B. gracilis*), and black grama (*B. eriopoda*) are both ecologically dominant and among the most valuable forage grasses of arid rangelands. Yet, genomic resources for the genus remain limited. We generated PacBio HiFi draft assemblies for *B. curtipendula* (653 Mb, N50 25.5 Mb), *B. gracilis* (808 Mb, N50 4.1 Mb), and *B. eriopoda* (1.67 Gb, N50 1.0 Mb). All assemblies showed >98.5% BUSCO completeness. Helixer predicted 83,877-215,536 gene models per assembly. However, syntenic copy numbers (2 in *B. curtipendula*, 3 in *B. gracilis*, and 6 in *B. eriopoda*) indicate that at least two species are polyploid, and the inflated gene model counts likely reflect homeologous copies and fragmented predictions rather than true gene numbers. panEDTA identified 25-40% transposable-element content, dominated by LTR retrotransposons. These assemblies provide foundational genomic references for *Bouteloua* and support comparative studies across PACMAD grasses.

## INTRODUCTION

Grasses (Poaceae) are ecologically and economically critical, with the PACMAD clade containing many C4 lineages adapted to arid and warm environments (Edwards and Still 2008). The tribe *Cynodonteae* (Chloridoideae) is one of the largest in the family, comprising ∼839 species in ∼93 genera, and includes agronomically important grasses such as finger millet (*Eleusine coracana*) and bermuda grass (*Cynodon dactylon*). It is further characterized by remarkable morphological diversity in inflorescence architecture (Pilatti et al. 2018). Within this clade, the Chloridoid genus *Bouteloua* is an ecologically important group of forage grasses comprising ∼60 species found exclusively in the Western Hemisphere, with a center of diversity in northern Mexico (Peterson et al. 2015). Commonly known as grama grasses, the genus is divided into two subgenera depending on whether the whole spikelet or the whole inflorescence branch falls from the mature plant. *Bouteloua* consists predominantly of perennial grasses, though a minority of species are annuals, with variable growth forms that may include bunch-forming, rhizomatous, or stoloniferous habits. Cytological studies indicate a common base chromosome number of x = 10, with widespread polyploidy contributing to its classification as a young polyploid complex (Gould 1979). Despite their ecological significance, genomic resources for *Bouteloua* remain limited compared to other Chloridoid grasses such as *Zoysia, Oropetium, Eleusine*, and *Eragrostis*.

Several *Bouteloua* species are ecologically dominant in North American grasslands. *B. curtipendula* (sideoats grama) ranges from Canada to South America and includes apomictic forms within a taxonomically complex group (Halbrook 2012); recent work has identified locally adapted ecotypes in the Chihuahuan Desert (Baez-Gonzalez et al. 2025). *B. gracilis* (blue grama) is common in the U.S. shortgrass steppe and Mexican highlands, showing phenotypic plasticity with molecular evidence for a Mexican origin and subsequent diversification northward (Avendaño-González et al. 2019). *B. eriopoda* (black grama) is a perennial, rhizomatous species of the Chihuahuan Desert, where recruitment patterns shape population persistence (Minnick and Coffin 1999). Cytological studies show extensive ploidy diversity: the *B. curtipendula* complex includes diploids and multiple polyploid forms (Siqueiros-Delgado et al. 2017), *B. gracilis* spans diploid to dodecaploid races (Snyder and Harlan 1953; Butterfield and Wood 2015), and *B. eriopoda* is predominantly diploid with occasional tetraploids (Streetman and Wright 1960).

Despite their ecological importance, flow-cytometric genome size data are lacking for these species, and genomic resources remain limited.

Here, we present high-quality draft genome assemblies and annotations for three *Bouteloua* species generated from PacBio HiFi long reads. Assemblies were evaluated with BUSCO, annotated with Helixer for gene models, and analyzed with panEDTA for transposable elements. These resources expand genomic coverage in the PACMAD clade and provide a foundation for investigating the genetic basis of ecologically and agriculturally important traits in *Bouteloua*, including adaptations to drought, extreme heat, and cold tolerance that shape their persistence in North American grasslands.

## METHODS & MATERIALS

### Plant Material, Sequencing, and Data Integrity

We sampled three *Bouteloua* species (*B. curtipendula, B. gracilis*, and *B. eriopoda*), each represented by a single individual grown from seed. *B. eriopoda* seed was sourced from Native American Seed (lot #2047932NM118; collected from New Mexico), *B. curtipendula* from Sheffield’s Seed Company (lot #1829746; collected from Texas), and *B. gracilis* from Great Basin Seed (lot #2681; collected from New Mexico). Plants were established in both a growth chamber (28 °C day/24 °C night) and a greenhouse (28 °C day/22 °C night) with a 14 h light/10 h dark photoperiod (supplemental lighting provided from 6:00-20:00, turned off when outdoor light exceeded 500 W m^−2^). Plants were grown in a soilless mix of peat, vermiculite, perlite, and turface, supplemented with limestone, calcium sulfate, and a 10-5-10 media mix fertilizer.

Fertilization followed a standard solution (15-5-15 Cal-Mag, 21-5-20 all-purpose, and Sprint 330 Fe chelate) applied at 150 ppm via dosatron three times per week, with water-only irrigation on other days. Young, healthy leaf tissue was collected from a single individual per species at the seedling stage, and high molecular weight DNA was extracted using the Macherey-Nagel NucleoBond® HMW DNA kit following the liquid nitrogen and mortar-and-pestle protocol (gravity-flow, buffers H1–H5). DNA was bound to the silica column, washed, and eluted in Buffer HE after isopropanol precipitation. Libraries were sheared using a MegaRuptor 3, size-selected on a PippinHT, and prepared with the PacBio SMRTbell Prep Kit 3.0. Sequencing was performed on the PacBio Revio platform using the 8 M SMRT Cell and Revio SPRQ Polymerase Kit, generating 35.97 Gb (*B. curtipendula*), 19.71 Gb (*B. gracilis*), and 24.81 Gb (*B. eriopoda*) of circular consensus reads. File integrity of the raw HiFi BAMs (and associated .bam.pbi indices) was verified by MD5 checksum comparison against provided hash manifests prior to downstream processing.

### Genome Assembly

For each species, we assembled genomes with hifiasm v0.25.0-r726 (64 threads) using default parameters for HiFi data with n-haps as 2 (Cheng et al. 2021). Assemblies were taken from the primary contig graph output; primary contig sequences were extracted from the GFA by selecting S-type records and writing them to FASTA.

### Assembly Evaluation

We quantified assembly size distributions with seqkit v0.15.0 and computed contig N50 (the contig length at which 50% of the assembly is contained in contigs of that size or longer) and L50 (the number of contigs needed to reach 50% of the assembly) using a custom length-sorting and cumulative-sum script (Shen et al. 2016). Assembly completeness was assessed with BUSCO v5.5.0 in genome mode against embryophyta_odb10 (64 threads) (Simão et al. 2015).

### Phylogeny Estimation

To identify syntenic regions, we used AnchorWave v1.2.6 (Song et al 2021) and the diploid outgroup *Oropetium thomeaum* (VanBuren et al. 2018), allowing each reference gene to participate in up to 12 alignment blocks. We sampled 100 gene regions with the modal copy number for the species, aligned these regions with MAFFT v7.525 with parameters –genafpair --maxiterate 1000 --adjustdirection (Katoh and Standley 2013), built gene trees with raxml-ng v1.2.0 using a GTR+G model (Kozlov et al. 2019), and combined them into a species tree with ASTRAL-PRO3 v1.23.3.6 (Zhang et al. 2025). To identify whole genome duplications on this species tree, we used GRAMPA v1.4.4 (Thomas et al. 2017) to reconcile gene duplications into a multiply-labeled species tree that indicates the position of polyploidy events.

### Gene Prediction and Basic Annotation

Structural gene annotation was performed with Helixer v0.3.5 using the land_plant model (Stiehler et al. 2021). For each assembly, Helixer produced GFF3 output, and coding sequence and peptide FASTA files. We enumerated gene counts from gene features and calculated CDS statistics (number of CDS features and average CDS length) from GFF3 coordinates. When required, CDS and protein sequences corresponding to annotated features were generated with gffread v0.9.12 against the species-specific genome FASTA (Pertea and Pertea 2020).

### Transposable element (TE) annotation and repeat metrics

Species-level TE libraries and genome annotations were generated with EDTA v2.1.0 using plant-agnostic settings --species others, --anno 1, with additional high-sensitivity re-runs where noted (Ou et al. 2024 Aug 1). For multi-genome summaries and harmonized outputs, we used panEDTA (wrapper around EDTA; inputs provided via local symlinks to avoid path issues).

From the resulting EDTA/panEDTA reports, we calculated the percentage of the genome masked overall and summarized the contribution of major superfamilies (e.g., LTR/Ty3, LTR/Copia, CACTA, Mutator, hAT, PIF/Harbinger, Helitron, LINE/SINE).

### Compute Environment and reproducibility

All analyses were executed on an HPC cluster under Linux using 64-128 CPU threads per job, where specified. Containerized tools (Helixer, EDTA/panEDTA) were run with Apptainer/Singularity to ensure version stability; native binaries were used for hifiasm, seqkit, gffread, and BUSCO. Exact software versions are reported above; commands were run with default parameters unless noted.

## RESULTS AND DISCUSSION

### Genome Assemblies

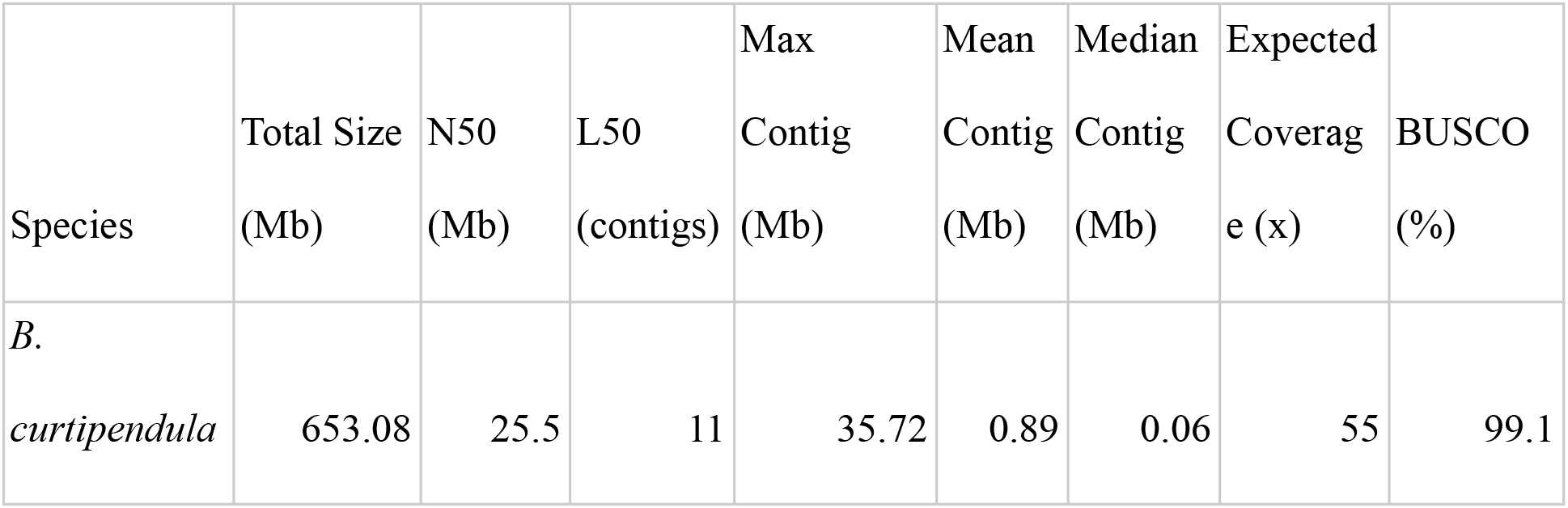
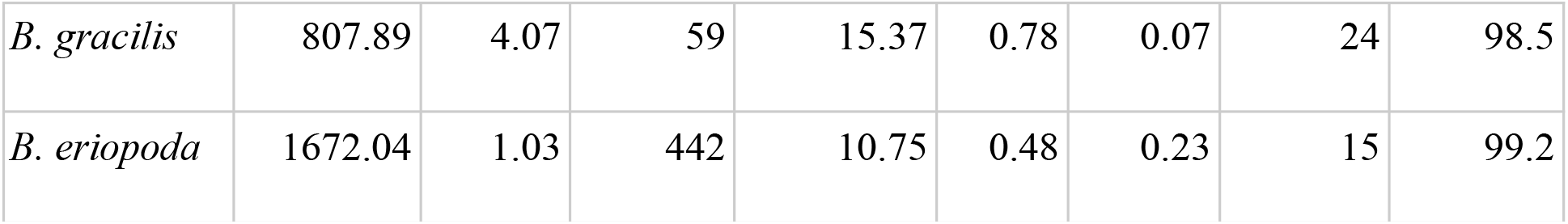

PacBio HiFi sequencing and hifiasm assembly produced high-quality draft genomes for *Bouteloua curtipendula, B. gracilis*, and *B. eriopoda* (Table 1). The assembly size of *B. curtipendula* was 653 Mb with an N50 of 25.5 Mb and L50 = 11, reflecting a highly contiguous genome. *B. gracilis* (808 Mb, N50 = 4.1 Mb, L50 = 59) and *B. eriopoda* (1.67 Gb, N50 = 1.0 Mb, L50 = 442) were more fragmented, with L50 increasing as assembly size increased. All assemblies exceeded 98.5% BUSCO completeness, indicating near-comprehensive gene space representation. Flow cytometry estimates of genome size of these species are rare in the literature, and are not connected to ploidy information. The assembly size of our *B. curtipendula* individual is similar to flow cytometry estimates of two accessions from Wisconsin (530 Mb and 644 Mb, (Bai et al. 2012), suggesting genome completeness. The only literature estimate of *B. curtipendula* genome size is 19 Gb (Galbraith et al. 1983), and is likely of higher ploidy than our sample, making comparisons difficult.

### Phylogeny Estimation

Syntenic copy numbers across species indicated that at least two of our sampled individuals are likely polyploid, although further cytological work would be needed to fully determine their ploidy levels. In *B. curtipendula*, most outgroup loci were represented by two syntenic copies, a pattern consistent with either divergent alleles in a diploid or two tetraploid subgenomes, each with low diversity. In *B. gracilis*, three syntenic copies were present for most loci, while *B. eriopoda* had six copies.

Polyploid species with divergent subgenome origins can be difficult to represent in a standard singly labeled species tree. To address this, we used GRAMPA to reconcile phylogenetic trees from 100 syntenic regions containing the most common 2/3/6 copies. The lowest-scoring multilabeled tree (reconciliation score 838; Figure 1D) suggests the *B. eriopoda* sample is an allopolyploid, with one parental subgenome closely related to *B. gracilis*.

**Figure 1:**
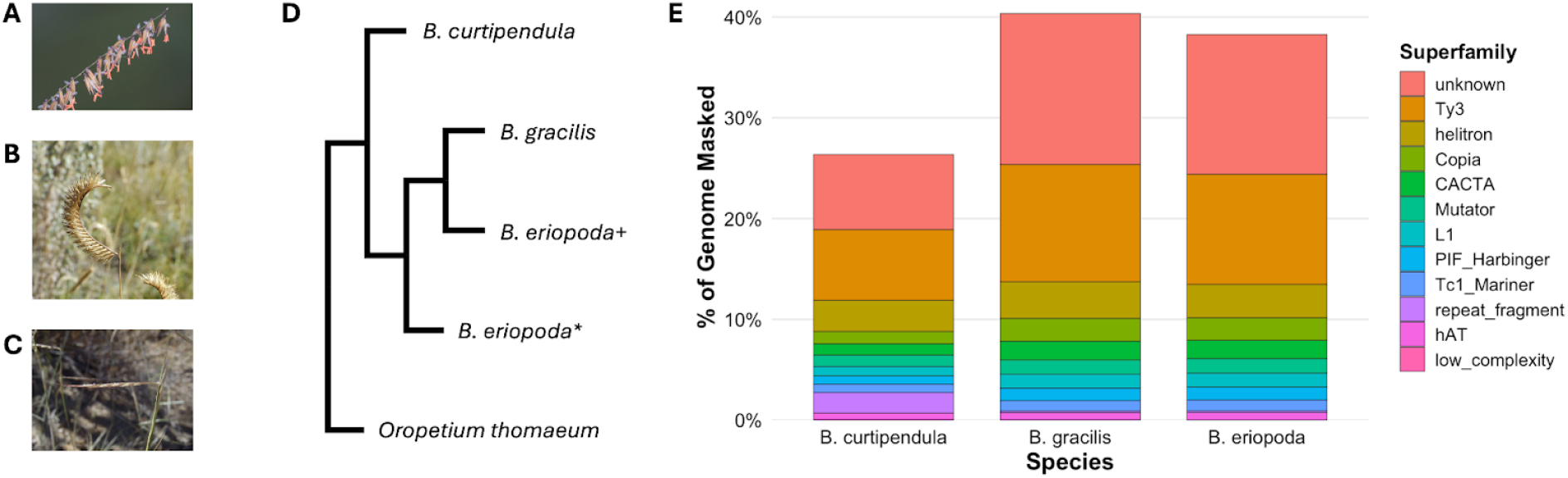
**A)** *B. curtipendula* “sideoats grama” Photo by BJ Stacey, distributed on iNaturalist under a CC BY-NC 4.0 license. **B)** *B. gracilis* “blue grama” Photo by Matthias Buck, distributed on iNaturalist under a CC BY-NC 4.0 license. **C)** *B. eriopoda* “black grama” Photo by Dan Beckman, distributed on iNaturalist under a CC BY-NC 4.0 license. **D)** Multilabeled species tree summarizing 100 loci. Separate subgenomes of *B. eriopoda* are indicated by “+” and “*.” **E)** Transposable element composition varied by superfamily in three *Bouteloua* species, showing higher Ty3 and unclassified content in *B. gracilis* and *B. eriopoda* relative to *B. curtipendula*.

### Gene Annotation

| Species | Genes | Avg CDS<br>length (bp) | TE %<br>Genome | Total Repeat Size<br>(Mb) | LTR<br>Retrotransposon<br>Content Size (Mb) |
| --- | --- | --- | --- | --- | --- |
| *B.*<br>*curtipendula* | 83877 | 1165.93 | 25.36 | 159.69 | 102.48 |
| *B. gracilis* | 104050 | 1128.92 | 40.3 | 317.46 | 233.51 |
| *B. eriopoda* | 215536 | 1149.15 | 38.63 | 628.67 | 452.05 |

Helixer predicted 83,877 genes in *B. curtipendula*, 104,050 in *B. gracilis*, and 215,536 in *B. eriopoda* (Table 2). Total gene numbers scaled with assembly size, suggesting that duplication and polyploidy have shaped *Bouteloua* genomes. These annotations represent the first gene catalogs for the genus and provide a foundation for future comparative and functional studies. However, these gene catalogs should be regarded as preliminary. The total gene counts (83,877–215,536) are considerably higher than expected for diploid grasses (∼30,000–40,000) (Devos 2010), reflecting a possible combination of polyploidy, allelic redundancy, and annotation fragmentation. Additionally, without transcriptomic evidence, some transposable elements and pseudogenes may be misclassified as coding sequences. Thus, while these annotations provide a useful starting point, refined pipelines that integrate transcriptomic and homology evidence will be required to generate a conservative set of nonredundant gene models for *Bouteloua*.

### Transposable Elements

LTR retrotransposons, particularly Ty3 and Copia, were the largest contributors. *B. gracilis* and *B. eriopoda* showed larger Ty3 expansions than *B. curtipendula*, consistent with their larger genome sizes. DNA transposons (CACTA, Mutator, hAT, PIF/Harbinger) and Helitrons contributed smaller fractions, while unclassified repeats were enriched in *B. gracilis* and *B. eriopoda*. Within Chloridoideae, these values fall between *Eragrostis tef* (∼27% repeats; ∼730 Mb genome) (Yg et al. 2016) and finger millet (*Eleusine coracana*; ∼50% repeats; ∼1.2-1.5 Gb) (Hittalmani et al. 2017).

Together, these resources create opportunities to investigate traits central to *Bouteloua*’s role in grassland ecosystems – from drought resilience to forage value – and to place *Bouteloua* more firmly in the comparative genomics landscape of PACMAD grasses. To date, only eight Chloridoideae species (*O. thomaeum, E. coracana, Z. japonica, Z. matrella, Z. matrella var. pacifica, E. nindensis, E. tef*, and *E. curvula*) have published nuclear genomes, highlighting that this subfamily remains comparatively undersampled (Iii 2025). Our assemblies, therefore, fill a key gap by providing new genomic resources to promote further study and conservation of a genus with a prominent role in North American drylands.

## DATA AVAILABILITY

The genome assemblies and annotations generated in this study are deposited on Figshare under the project High-Quality Draft Genome Assemblies and Comparative Genomics of Three *Bouteloua* Species (Poaceae: Chloridoideae). The dataset for *Bouteloua curtipendula* is available at https://doi.org/10.6084/m9.figshare.30311320, the dataset for *Bouteloua eriopoda* is available at https://doi.org/10.6084/m9.figshare.30311338, and the dataset for *Bouteloua gracilis* is available at https://doi.org/10.6084/m9.figshare.30311368. Each dataset includes the draft genome assembly, structural and functional annotations, and associated metadata. The inputs to analyses, code to reproduce tables and plots, and summary tables are available in the GitHub repository https://github.com/AGanesan12/Bouteloua-Assembly-and-Annotation. All raw

PacBio HiFi sequencing reads, assembled genomes, and annotation files generated in this study are also available from the corresponding author upon request.

## ACKNOWLEDGEMENTS

We thank the Buckler Lab, especially Thuy La, Nick Lepak, Ana Berthel, Zack Miller, Sheng-Kai Hsu, and Mohammed El-Walid, for support and advice.

ChatGPT (OpenAI GPT-5, Sept 2025) was used to refine wording in non-technical sections; all analyses and interpretations were performed by the authors.

## FUNDING

This work was supported by USDA-ARS Project Number 8062-21000-052-004-A, the Institute for Genomic Diversity at Cornell University, and the Buckler Lab computing cluster (BioHPC).

E.S.B. is supported by the USDA-ARS (ARS project number 8062-21000-052-000-D).

## CONFLICTS OF INTEREST

The authors declare that there is no conflict of interest.

## Notes

https://doi.org/10.6084/m9.figshare.30311320

https://doi.org/10.6084/m9.figshare.30311338

https://doi.org/10.6084/m9.figshare.30311368

### Competing Interest Statement

The authors have declared no competing interest.

## REFERENCES

Avendaño-González M, Morales-Domínguez JF, Siqueiros-Delgado ME. 2019. Genetic structure, phylogeography, and migration routes of Bouteloua gracilis (Kunth) Lag. ex Griffiths (Poaceae:Chloridoideae). Mol Phylogenet Evol. 134:50–60. 10.1016/j.ympev.2019.01.005

Baez-Gonzalez AD et al. 2025. A GIS Approach to Modeling the Ecological Niche of an Ecotype of Bouteloua curtipendula (Michx.) Torr. in Mexican Grasslands. Plants. 14(14):2090. 10.3390/plants14142090

Bai C, Alverson WS, Follansbee A, Waller DM. 2012. New reports of nuclear DNA content for 407 vascular plant taxa from the United States. Ann Bot. 110(8):1623–1629. 10.1093/aob/mcs222

Butterfield BJ, Wood TE. 2015. Local climate and cultivation, but not ploidy, predict functional trait variation in Bouteloua gracilis (Poaceae). Plant Ecol. 216(10):1341–1349. 10.1007/s11258-015-0510-8

Cheng H et al. 2021. Haplotype-resolved de novo assembly using phased assembly graphs with hifiasm. Nat Methods. 18(2):170–175. 10.1038/s41592-020-01056-5

Devos KM. 2010. Grass genome organization and evolution. Curr Opin Plant Biol. 13(2):139–145. 10.1016/j.pbi.2009.12.005

Edwards EJ, Still CJ. 2008. Climate, phylogeny and the ecological distribution of C4 grasses. Ecol Lett. 11(3):266–276. 10.1111/j.1461-0248.2007.01144.x

Galbraith DW et al. 1983. Rapid Flow Cytometric Analysis of the Cell Cycle in Intact Plant Tissues. Science. 220(4601):1049–1051. 10.1126/science.220.4601.1049

Gould FW. 1979. The Genus Bouteloua (Poaceae). Ann Mo Bot Gard. 66(3):348–416. 10.2307/2398834

Halbrook AK. 2012. BOUTELOUA CURTIPENDULA (POACEAE): REPRODUCTIVE BIOLOGY, PHENOTYPIC PLASTICITY, AND THE ORIGINS OF AN APOMICTIC SPECIES COMPLEX.

Hittalmani S et al. 2017. Genome and Transcriptome sequence of Finger millet (Eleusine coracana (L.) Gaertn.) provides insights into drought tolerance and nutraceutical properties. BMC Genomics. 18:465. 10.1186/s12864-017-3850-z

Iii GPWG. 2025. A nuclear phylogenomic tree of grasses (Poaceae) recovers current classification despite gene tree incongruence. New Phytol. 245(2):818–834. 10.1111/nph.20263

Katoh K, Standley DM. 2013. MAFFT Multiple Sequence Alignment Software Version 7: Improvements in Performance and Usability. Mol Biol Evol. 30(4):772–780. 10.1093/molbev/mst010

Kozlov AM et al. 2019. RAxML-NG: a fast, scalable and user-friendly tool for maximum likelihood phylogenetic inference. Bioinformatics. 35(21):4453–4455. 10.1093/bioinformatics/btz305

Minnick TJ, Coffin DP. 1999. Geographic patterns of simulated establishment of two Bouteloua species: implications for distributions of dominants and ecotones. J Veg Sci. 10(3):343–356. 10.2307/3237063

Ou S et al. 2024. Differences in activity and stability drive transposable element variation in tropical and temperate maize. [accessed 2025 Sept 13]. https://genome.cshlp.org/content/34/8/1140. 10.1101/gr.278131.123

Pertea G, Pertea M. 2020. GFF Utilities: GffRead and GffCompare. F1000Research. 9:ISCB Comm J-304. 10.12688/f1000research.23297.2

Peterson PM, Romaschenko K, Arrieta YH. 2015. Phylogeny and subgeneric classification of Bouteloua with a new species, B. herrera-arrietae (Poaceae: Chloridoideae: Cynodonteae: Boutelouinae). J Syst Evol. 53(4):351–366. 10.1111/jse.12159

Pilatti V et al. 2018. Diversity, systematics, and evolution of Cynodonteae inflorescences (Chloridoideae – Poaceae). Syst Biodivers. 16(3):245–259. 10.1080/14772000.2017.1392371

Shen W, Le S, Li Y, Hu F. 2016. SeqKit: A Cross-Platform and Ultrafast Toolkit for FASTA/Q File Manipulation. [accessed 2025 Sept 15]. https://journals.plos.org/plosone/article?id=10.1371/journal.pone.0163962. 10.1371/journal.pone.0163962

Simão FA et al. 2015. BUSCO: assessing genome assembly and annotation completeness with single-copy orthologs. Bioinformatics. 31(19):3210–3212. 10.1093/bioinformatics/btv351

Siqueiros-Delgado ME, Fisher AE, Columbus JT. 2017. Polyploidy as a Factor in the Evolution of the Bouteloua curtipendula Complex (Poaceae: Chloridoideae). Syst Bot. 42(3):432–448. 10.1600/036364417X696159

Snyder LA, Harlan JR. 1953. A Cytological Study of Bouteloua gracilis from Western Texas and Eastern New Mexico. Am J Bot. 40(9):702–707. 10.2307/2439685

Stiehler F et al. 2021. Helixer: cross-species gene annotation of large eukaryotic genomes using deep learning. Bioinformatics. 36(22–23):5291–5298. 10.1093/bioinformatics/btaa1044

Streetman LJ, Wright N. 1960. A Cytological Study of Black Gramagrass BouTELoua Eriopoda. Am J Bot. 47(9):786–793. 10.1002/j.1537-2197.1960.tb07166.x

Thomas GWC, Ather SH, Hahn MW. 2017. Gene-Tree Reconciliation with MUL-Trees to Resolve Polyploidy Events. Syst Biol. 66(6):1007–1018. 10.1093/sysbio/syx044

VanBuren R, Wai CM, Keilwagen J, Pardo J. 2018. A chromosome-scale assembly of the model desiccation tolerant grass Oropetium thomaeum. Plant Direct. 2(11):e00096. 10.1002/pld3.96

Yg G E B, Me P A Z. 2016. Identification and characterization of abundant repetitive sequences in Eragrostis tef cv. Enatite genome. PubMed. [published online ahead of print] [accessed 2025 Sept 13]. https://pubmed.ncbi.nlm.nih.gov/26833063/

Zhang C, Nielsen R, Mirarab S. 2025. ASTER: A Package for Large-Scale Phylogenomic Reconstructions. Mol Biol Evol. 42(8):msaf172. 10.1093/molbev/msaf172

